# Paradoxical relationship between speed and accuracy in olfactory figure-background segregation

**DOI:** 10.1101/2021.03.10.434790

**Authors:** Lior Lebovich, Michael Yunerman, Viviana Scaiewicz, Yonatan Loewenstein, Dan Rokni

## Abstract

In natural settings, many stimuli impinge on our sensory organs simultaneously. Parsing these sensory stimuli into perceptual objects is a fundamental task faced by all sensory systems. Similar to other sensory modalities, increased odor backgrounds decrease the detectability of target odors by the olfactory system. The mechanisms by which background odors interfere with the detection and identification of target odors are unknown. Here we utilized the framework of the Drift Diffusion Model (DDM) to consider possible interference mechanisms in an odor detection task. We consider effects of background odors on both signal and noise in the decision-making dynamics, and show that these produce different predictions about decision accuracy and speed. To test these predictions, we trained mice to detect target odors that are embedded in random background mixtures in a two-alternative choice task. Trial duration was independent of behavioral reaction times to avoid motivating rapid responses. We found that the behavioral data is most consistent with background odors acting by adding noise to the decision-making dynamics. The added noise decreases the correct rate, but also decreases decision times, thereby creating a paradoxical relationship between speed and accuracy of target detection, where mice make faster and less accurate decisions in the presence of background odors.

## Introduction

Natural scenes are cluttered and sensory systems must be able to parse these scenes to detect and identify individual objects such as food, a mate, or a predator. Scene segmentation is critical for all sensory modalities yet the neural mechanisms that underlie this feat are not very well understood. While these mechanisms have been extensively studied in the visual and auditory modalities (Crouzet and Serre, 2011; Elhilali and Shamma, 2008; Lamme, 1995; McDermott, 2009; Micheyl and Oxenham, 2010; Teki et al., 2011; Wolfson and Landy, 1998), very little is known about scene segmentation of olfactory scenes (Grabska-Barwinska et al., 2013; Li and Hertz, 2000; Mathis et al., 2016; Rokni et al., 2014).

We have previously developed a psychophysical paradigm for testing detection of target odors against background mixtures in mice (Rokni et al., 2014). Utilizing this paradigm, we found that the ability to report whether a target odor is present decreases when the number of background odors is increased. Additionally, we found that the overlap between the glomerular activation patterns that represent the target, and those that represent the background odors, was inversely correlated with success rate. This result suggested that background odors may interfere with target-odor detection already at the level of olfactory receptors. Indeed, many studies demonstrated non-linear interactions between odorants activating the same receptor (Duchamp-Viret et al., 2003; Firestein and Shepherd, 1992; Kurahashi et al., 1994; Oka et al., 2004; Pfister et al., 2020; Rospars et al., 2008; Singh et al., 2019; Takeuchi et al., 2009), and recent studies suggest such interactions are widespread (Reddy et al., 2018; Xu et al., 2020; Zak et al., 2020). However, it is currently unclear how these interactions affect target odor detection and in what way do they interfere with the target signal.

The drift-diffusion model (DDM) has been instrumental in providing insights into the effects of sensory signals on success rate and reaction times in sensory-guided tasks (Brody and Hanks, 2016; Gold and Shadlen, 2007; Krajbich and Rangel, 2011; Laming, 1968; Ratcliff, 1978; Ratcliff and McKoon, 2007). According to the DDM, the decision process is determined by a noisy decision variable, which reflects the difference between the accumulated evidence in favor of choosing each of the two alternatives (in this case, the presence *vs* the absence of the target odor). The decision is made once the decision variable reaches for the first time one of two thresholds, ±*θ*, where *θ* > 0. If the decision variable first reaches one of them (*θ*), the decision is to report that the target is present whereas if it first reaches the other (−*θ*), it reports that the target is absent.

The DDM is characterized by several parameters: the first parameter is the starting point m *θ* (−1 < *m* < 1), which denotes an a-priori, evidence-independent preference towards one of the outcomes. *m* = 0 indicates an unbiased starting point. An a-priori preference towards reporting that the target is present or absent manifests as a positive or negative value of *m*, respectively. The second parameter is the threshold *θ* > 0. The larger the value of *θ*, the more evidence is required in order to reach a decision. Third is the average rate of evidence accumulation in the presence and in the absence of the target, which we denote by *A*_+_ and *A*_−_, respectively. We assume that in absence of any odors, 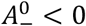, which biases the decision variable in the direction of the −*θ* threshold and the decision that the target is absent. By contrast, when the target odor is presented to the animal, 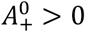 and as a result, the decision variable drifts (on average) towards the *θ* threshold. Finally, the decision variable also accumulates white noise in the decision process, and the magnitude of this noise is denoted by 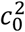 (Bogacz et al., 2006; Heekeren et al., 2008; Ratcliff, 1978; Ratcliff and McKoon, 2007; Ratcliff et al., 1999, 2003).

In this framework, there are at least three different mechanisms by which background odors can interfere with target odor detection. First, they may reduce the evidence in favor of the target, thereby increasing the probability that subjects will miss the target. Second, they may increase the evidence in favor of the target, thus generating a false target-like signal even when the target is absent, resulting in an increased false-alarm rate. Finally, they may act by increasing the noise in the decision-making process. Here we analyze the predictions of the DDM for each of these modes of interference and compare them to experimental observations.

## Results

We trained mice to detect target odorants against background mixtures. The task is similar to the one previously used (Rokni et al., 2014), but modified from a go/no go to a two-alternative choice reaction time design to allow analysis of reaction times in all trials. In each trial, mice were presented with a pseudorandom odorant mixture. Mixtures either included one of two target odorants (“target-on” trials), or neither of the target odorants (“target-off” trials). Mice were rewarded with a water drop if they correctly reported whether the mixture contained a target odorant by licking to the right in target-on trials and licking to the left in target-off trials (Figure 1). To avoid motivating rapid responses, trial duration was independent of response time and was 10 seconds for correct trials, and 15 seconds for incorrect trials. The rate of rewards was therefore dependent on the correct rate, but not the speed of behavioral responses. We trained 6 mice and collected an average of 16790 trials per mouse (range 12000-23000).

**Figure 1:**
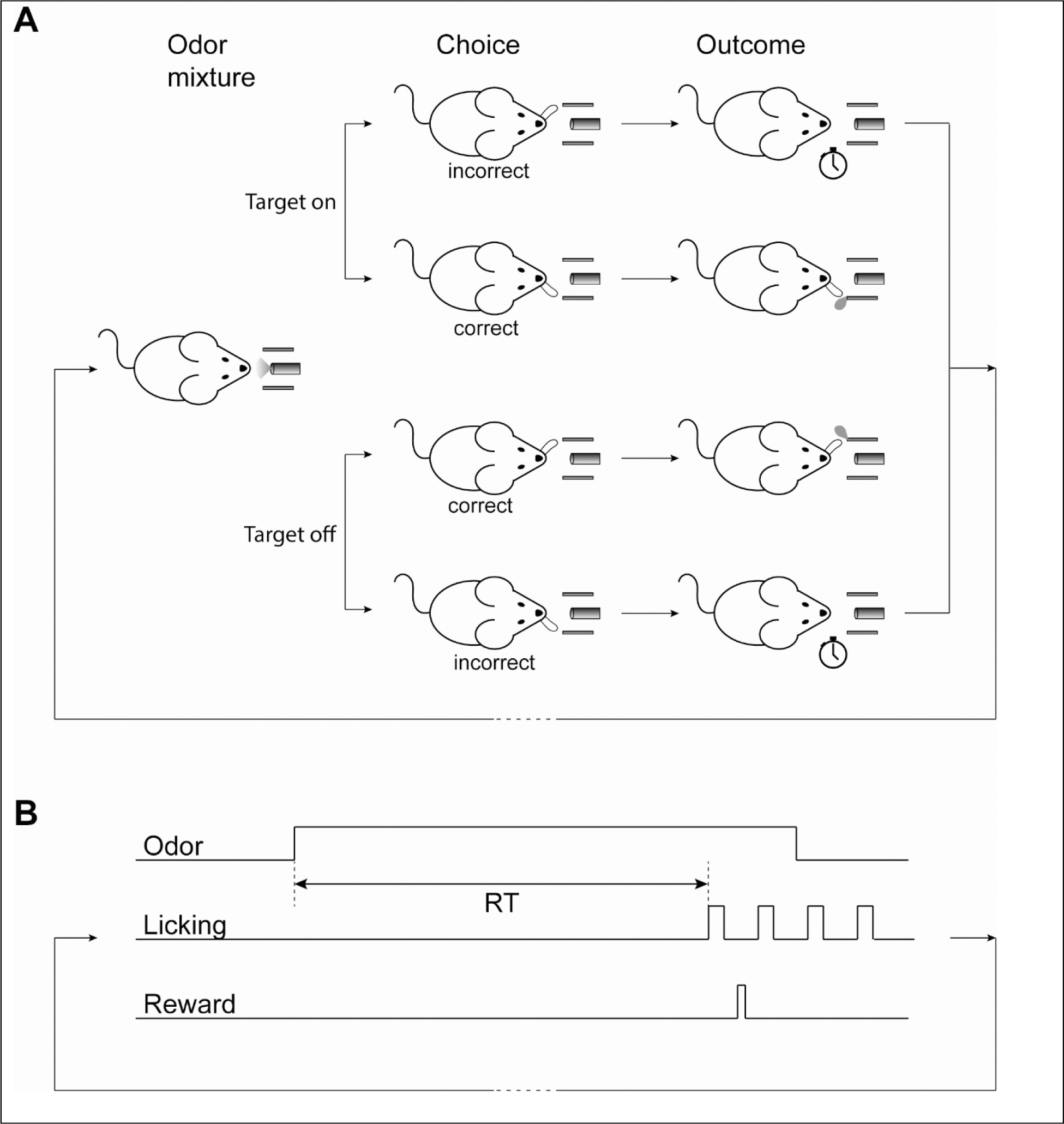
Behavioral paradigm. **A**. Mice were trained on a target detection with background task that was set as a two-alternative choice reaction time task. In each trial, mice were presented with an odorant mixture and were required to report the presence or absence of a target odorant by licking the right or left lick ports, respectively. Correct licks were rewarded with a water drop and incorrect licks were punished with a timeout. **B**. The temporal structure of a trial. Following odor onset, mice reported their decision by licking. The time between odor onset and the first lick was taken as the trial’s reaction time.

We considered three different mechanisms by which background odorants can affect decisions in the framework of the DDM and studied their behavioral predictions:

(1) False signal: Individual odors activate a large number of olfactory receptors. It is most likely that many of the target-activated receptors are also activated by the background odors (Rokni et al., 2014). Therefore, it is possible that the addition of background odorants to the odor mixture will manifest as an *increase* in the drift rates both in the presence of a target odor (*A*_+_) and in its absence (*A*_−_). The larger the number of background odors, the larger the drift rates would be.

(2) Signal reduction: Background odors, however, also activate olfactory receptors that are not activated by the target odors. Since inhibitory interactions are common in second and third order olfactory brain regions (Arevian et al., 2008; Aungst et al., 2003; Economo et al., 2016; Franks et al., 2011; Poo and Isaacson, 2009, 2011; Suzuki and Bekkers, 2012; Urban and Sakmann, 2002), addition of background odorants may inhibit target-associated signals. Such interactions are predicted to manifest as a *decrease* in the drift rates both in the presence of a target odor (*A*_+_) and in its absence (*A*_−_). The magnitude of the decrease would be a monotonic function of the number of background odors.

(3) Noise boost: Background odors may affect the dynamics of the target odor responses without a consistent effect on mean response. In that case the addition of background odorants could manifest as an increase in the noise associated with the decision process, *c*^2^.

The behavioral predictions of the three hypotheses are depicted in figure 2A-C (also see Eq. 2, 3, and 5, in the Methods). The “false signal” hypothesis predicts that an increase in the number of background odors will result in an increase in the target detection probability, but will also manifest in an increase in the number of false detections (Fig. 2A). In other words, the probability that the subject would report that the target is present *p* would increase with the number of background odorants, both when the target is present (blue) and when it is absent (red). In contrast, the “signal reduction” hypothesis makes the opposite prediction -the probabilities of both true and false detections will decrease with the number of background odors (Fig. 2B). Finally, the “noise boost” hypothesis predicts that with the increase in the number of background odorants, choice behavior would contract to a stimulus-independent probability ((*m* + 1)/2) resulting in both decrease in the probability of target detection and an increase in the probability of false detection (Fig. 2C). Importantly, these predictions are generic to this model and do not depend on a specific choice of parameters (as long as 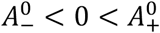). The experimental relationship between the number of background odorants and *p* for both trial types is depicted in figure 2D (thin lines, single subjects, thick lines, average over subjects). *P* decreased with the number of background odorants in target-on trials, and increased with the number of background odorants in target-off trials. This dependence matches the prediction of the “noise boost” hypothesis and is inconsistent with the “false signal” and “signal reduction” hypotheses.

**Figure 2:**
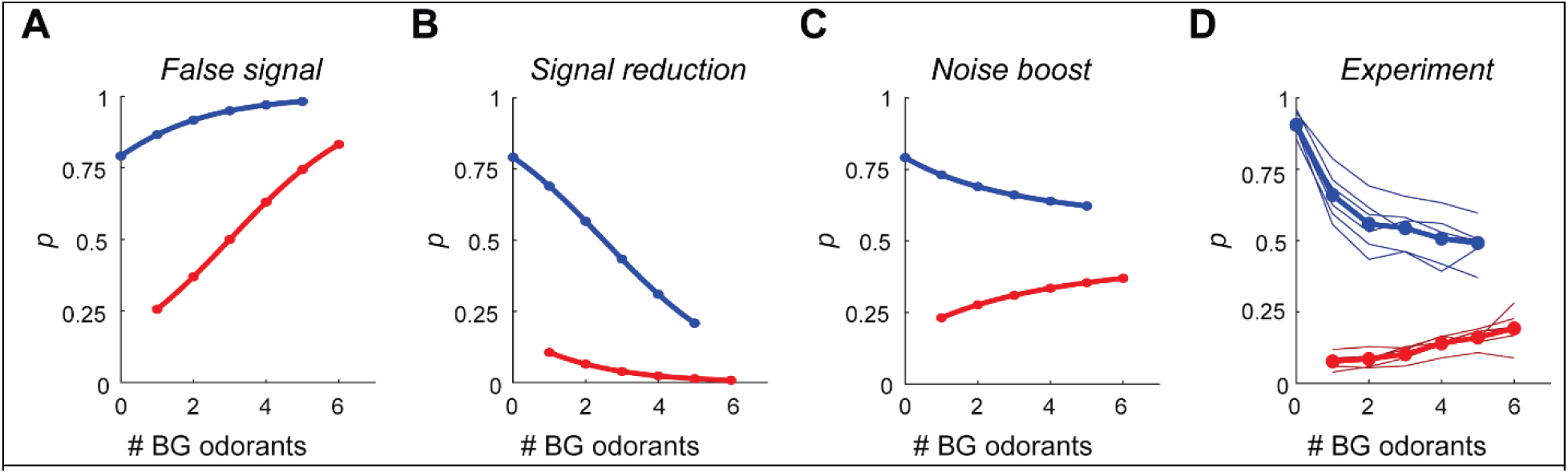
The effect of background odorants on decision probabilities. **A-C**. The probability of reporting that the target odorant is present (*p*) as a function of the number of background odorants, for the three interference hypotheses: false signal (**A**), signal reduction (**B**), and noise boost (**C**). Blue – “target on” trials, red – “target off” trials. Baseline parameter values are 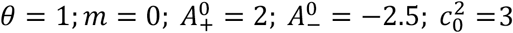. The drift rate under the false signal, the signal reduction and the noise boost mechanisms are: 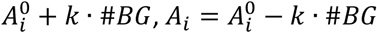 and 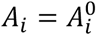, respectively, where *i* ∈ {+ −} and #*BG* is the number of background odorants in the mixture and *k*= 1. The diffusion is 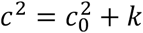 under the noise boost mechanism and 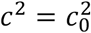 in the false signal and signal reduction mechanisms. **D**. The fraction of trials that mice reported that the target is present as a function of the number of background odorants. Thin lines show data from individual mice. Thick lines show the mean across mice. Colors as in A-C.

The “noise boost” hypothesis predicts that the average decision time would decrease with the number of background distractors both in target-on and in target-off trials (Fig. 3A). To gain insight to this prediction, consider the limit in which the variance of the noise in the decision process is very large. Because the decision variable integrates this noise, the magnitude of the decision variable will very quickly become very large, and will cross one of the decision thresholds. This prediction is counter-intuitive because it suggests that decisions in the more difficult trials -trials associated with more background odorants and hence more errors -will be made faster. To test this prediction, we measured the speed at which behavioral responses were made (reaction time). Although reaction times include other components beyond decision times, they are useful correlates of decision times. In agreement with the “noise boost” hypothesis, we found that reaction times decreased with the number of background odorants (Figure 3B). This finding is in contrast to a previous study that found a very small effect of background odorants on reaction times, yet that study used a go no go paradigm and was based on a much smaller dataset (Rokni et al., 2014). Taken together, our results suggest that the effects of background odorants on target detection are best explained by a mechanism in which adding background manifests as added noise in the decision making dynamics.

**Figure 3:**
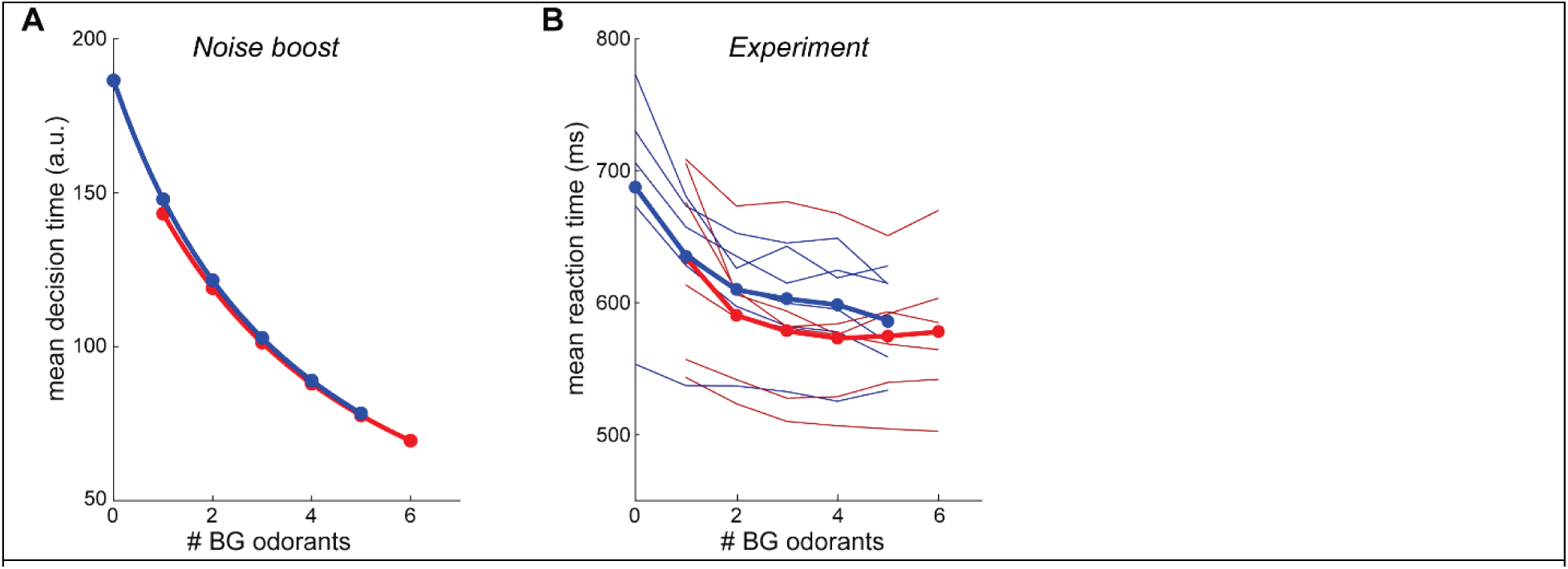
The effect of background odorants on decision times. **A**. Mean decision time under the “noise-boost” mechanism as a function of the number of background odorants. Blue – “target on” trials, red – “target off” trials. **B**. The mean mouse reaction time as a function of the number of background odorants. Thin lines show data from individual mice, and thick lines show the mean across mice. Colors as in A.

## Discussion

We used the DDM in order to understand how background odors affect the detectability of target odors. We considered three alternative mechanisms by which background odors may act: inducing target-like signal, reducing the target signal or increasing noise. The finding that mice make faster and less accurate decisions on trials with more background is only consistent with the increased noise hypothesis.

While our results are general and do not depend on the assumption of specific model parameters, several limitations of our analysis are worthwhile noting. First, we assumed that the background odors act in a singular fashion, either affecting the drift, or affecting the noise. It is possible however, that background odors interfere with both the drift and the noise. Second, our analysis implicitly assumed that all background odors have the same effect, ignoring their identity, as well as the identity of the target odors. Chemical and representational overlap between the target and background odorants were shown to determine background potency (Rokni et al., 2014). The magnitude of noise added by background odorants, may therefore be odorant dependent. Additionally, it is possible that some odorants affect the drift. A larger behavioral dataset may allow further separation of trials based on specific target-background combinations. Finally, our analysis is fully based on a specific theoretical framework – the DDM.

Importantly, our study is, to our knowledge, the first to find an inverse relationship between trial difficulty and reaction time in an odor-guided task. Previous studies used odor discrimination tasks to analyze the relationship between speed and accuracy (Abraham et al., 2004; Mendonça et al., 2020; Rinberg et al., 2006; Uchida and Mainen, 2003; Zariwala et al., 2013). In such tasks the difficulty is presumed to be related to the signal (odor similarity), and yet contrasting results regarding the effect of discrimination difficulty on reaction times were reported. It is not entirely clear what gave rise to the different results, yet it is possible that the difficulty in some of the tasks that were used, was accurately representing the discrimination boundaries rather than sensory stimulus itself (Zariwala et al., 2013). Additionally, a recent study indicated that the learning process may also contribute to the relationship between task difficulty, success rate and reaction times (Mendonça et al., 2020).

What do our results teach us about cross-odorant interactions? The overlapping nature of sensory representations in the olfactory epithelium provides opportunity for many non-linear odorant interactions (Araneda et al., 2000; Malnic et al., 1999; Meister and Bonhoeffer, 2001; Rubin and Katz, 1999; Soucy et al., 2009). Interaction between odorants have been studied mostly at the level of sensory neurons (Duchamp-Viret et al., 2003; Firestein and Shepherd, 1992; Kurahashi et al., 1994; Oka et al., 2004; Pfister et al., 2020; Rospars et al., 2008; Singh et al., 2019; Takeuchi et al., 2009; Xu et al., 2020; Zak et al., 2020) but also, a bit more anecdotally, in the olfactory bulb and cortex (Davison and Katz, 2007; Fletcher, 2011; Giraudet et al., 2002; Gupta et al., 2015; Kadohisa and Wilson, 2006; Lei et al., 2006; Penker et al., 2020; Stettler and Axel, 2009; Wilson, 2003; Yoshida and Mori, 2007). These studies analyzed the amplitudes of responses to mixtures of odorants and found that these are typically sublinear. The diffusion term in the DDM represents noise that evolves with time within a trial. The finding that background odorants act as noise in the decision making process may therefore suggest that adding background odorants changes not only the amplitude of odor responses but also response dynamics. It will be interesting to analyze the dependence of response dynamics on background in olfactory brain regions.

Background segmentation is a task that is faced by all sensory systems. Is our finding that reaction times decrease with increased background specific to the olfactory system? Although background segmentation has been rather extensively studied in both vision and audition, there are, as far as we know, no studies that immediately compare to ours. When detecting Gabor patterns in noise, the number of eye fixations required for detection increases with various parameters of the stimulus, yet the effects of varying the noise on reaction times have not been examined (Najemnik and Geisler, 2005). Reaction times in tone in noise detection in the auditory system increase when the signal to noise ratio is decreased (either by reducing the signal or enhancing the noise), but the noise in these tasks was continuously present throughout the session and did not obey the trial structure (Dylla et al., 2013; Kemp, 1984). Future experiments that vary the background with a trial structure may promote comparison of the effects of background on reaction times in the different sensory systems.

## Methods

All experimental procedures were performed using approved protocols in accordance with institutional (Hebrew University IACUC) and national guidelines.

### Behavior

#### Subjects and Surgery

6 c57bl6 adult male mice (Envigo) were trained on the behavioral task. Mice were first anesthetized (Ketamine/Xylazine 100 and 10 mg/kg, respectively) and a metal plate was attached to their skull with dental acrylic for subsequent head restraining. Mice were then maintained in a reversed light/dark cycle facility. All behavioral training and testing was done during the subjective night time.

#### Apparatus

The behavioral apparatus was located inside a sound attenuating box (Med Associates, VT USA) and consisted of a head restraining device, an odor delivery system, a dual lick detector and a water delivery system. Odor delivery, monitoring of licking and water rewards were controlled using computer interface hardware (National Instruments) and custom software written in LabVIEW. The mouse was continuously monitored using a CCD camera during behavior sessions under red illumination.

#### Odor presentation

Odorant mixtures were presented using a custom-made odor machine as described previously (Rokni et al., 2014). The odor machine was designed to supply constant flow (1.5 liters/minute) and have the concentrations of the different odorants independent of each other. The odor machine was composed of 8 odor modules. Each module was made of two glass tubes, one containing the odor and solvent and the other containing only the solvent. A 3-way valve (Lee Company, USA) diverted an input flow of filtered air to either the odor tube or the solvent tube, and the output of both tubes was merged to form the module output. This design ensured that each module contributed a constant amount of flow at any time. Input flow to the modules and output flow from the modules were made of ultra-chemical-resistant Tygon/PVC tubing connected in symmetric pair-wise bifurcations to ensure equal flow on all modules. All odorants were diluted to 10% v/v in diethyl phthalate (Sigma Aldrich, CAS 84-66-2) in the tubes and then further diluted in gas phase by the flow of other modules 8 fold. The odorant mixture was carried from the point of final odorant convergence to the odor port through a 1-meter-long tubing with an inner diameter of 1/16 inch to allow mixing while minimizing the latency from valve opening to odor presentation at the mouse’s nostrils. The time delay between valve opening and odor delivery (200 ms), was measured using a photoionization detector (miniPID, Aurora Scietific), and was subtracted from all reaction time values.

#### Odor set

All odorants were obtained from Sigma Aldrich. The odorants were (CAS number in parenthesis): Ethyl propionate (105-37-3), Isoamyl tiglate (41519-18-0), Ethyl tiglate (5837-78-5), 2-Ethylhexanal (123-05-7), Propyl acetate (109-60-4), Isobutyl propionate (540-42-1), Ethyl valerate (539-82-2), Phenethyl tiglate (55719-85-2). The target odorant pairs and the mice trained to detect each pair are as follows: Ethyl propionate and Isoamyl tiglate – mice 1&2. Ethyl tiglate and 2-Ethylhexanal – mice 3&4. Propyl acetate and Isobutyl propionate – mice 5&6.

#### Behavioral training and testing

Following surgery mice were allowed one week of recovery and were then water restricted. Mice were then acclimatized to the behavioral apparatus for at least 2 days in which they were allowed 30 minutes of free exploration and free water at the apparatus. This was followed by 2 days in which mice were head-restrained and were rewarded with a water drop for licking the water spout. On the fifth day of water restriction mice began training on the task. No randomized process was used to assign target odorants to each mouse. A mixture was presented for 2 seconds every 10 seconds and mice had to lick to the right if the mixture included one of the two target odorants (Ton) and to the left if it did not (Toff). Mice had to respond within a 2.8 second period (starting 200 ms after odor onset). Correct licks were rewarded with an 8µL water drop, and incorrect licks were punished by a 5 second time-out. Rejections were not punished and not rewarded. Training began with easy sessions and as mice reached 70% performance over a whole session, session difficulty was adjusted. The difficulty of the task was controlled by varying the distribution of the number of components in the mixture using the following equation:

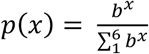

where p(x) is the probability of x components (ranging from 1 to 6). The parameter b was first set to 0.1 and was then raised sequentially through 0.25, 0.5, 0.75 reaching a maximal value of 1. Data was collected only in session with b>=0.5. Mice performed one session per day with an average of 250 ± 20 trials per session (lasting typically about an hour), and each mouse contributed between 60 and 91 sessions to the data set (72 ± 12 mean ± SD, n=6 mice). Mice often had a bias towards one side (initially often licking to only one side). To eliminate this bias as much as possible, stimulus statistics (Ton and Toff) were made dependent on recent choices. The chance of the next trial being Ton was one minus the proportion of licks to the right in the last 5 trials. Thus if a mouse preferred licking to the left, he would get more trials in which the correct response is to lick to the right. Chance level performance (the performance if mice just guess and ignore the stimulus) is not easily defined here because the bias is not known, however it is always less than 50%.

#### Data analysis

All analysis was performed using custom written code in Matlab. Sessions with complexity below 0.5 were considered training sessions and were not analyzed (except for learning curves). Occasional testing sessions in which performance dropped below 70% were also not analyzed (mostly after breaks in training). All reported parameters were calculated for each mouse separately. To remove variation in reaction times that stem from variable positioning of the water spouts, the reaction times in each session were normalized to the sessions mean. All reaction times were then multiplied by the mean reaction time across all sessions to provide meaningful units.

### Modeling

According to the DDM, the decision is made once a decision variable *x*, which reflects the difference between the accumulated evidence in favor of choosing each of the two alternatives, reaches one of two thresholds for the first time, ±*θ*, where *θ* > 0. Formally, the dynamics within a trial is given by *dx* / *dt* = *A* + ξ where *A* is the drift rate, *t* is time and ξ denotes white noise such that E[ξ(*t*)] = 0 and E[ξ (*t*) ξ (*t* ′)] = *c*^2^ δ (*t* − *t* ′). The initial state of the decision variable is given by *x*(*t* = 0) = *mθ* (*m* ∈ (−1 1)). In this model, the probability that the decision variable will reach +*θ* first, *p*, as a function of the model parameters is given by (Bogacz et al., 2006; Srivastava et al., 2016):

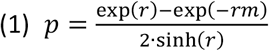

where *r* = 2*Aθ c*^2^ ≠ 0. We denote by +*θ* and −*θ* the decision thresholds associated with “target present” and “target absent” decisions and by *A*_+_ and *A*_−_ the drift rates associated with trials in which the target odor is present (“target on” trials) and absent (“target off” trials), respectively, and assume that in the absence of background odors, 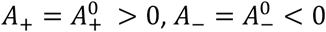 and 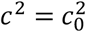.

To study the effect of changing the drift rates or the variance of the noise on *p*, we compute the partial derivative of *p* with respect to these variables:

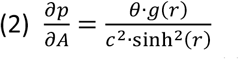

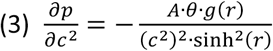

where: *g* (*r*) =exp (− *rm*) · [cos (*r*) + *m* · sinh (*r*)] − 1.

We note that *g* (*r*), in the domain *D* = {(*r m*) | (−1 < *r.m* < 1) ^ *m* ∈ ℝ}, is bounded from below by inf {*g* (*r*) | (*r*, *m*) ϵ*D* = 0. This is because the first derivative of *g* (*r*) in *D, ∂g / ∂r* = *e* ^−*rm*^ · (1 − *m*^2^) · sinh (*r*) = 0 only for *r* = 0, the value of the second derivative of *g* (*r*) in *D* at *r* = 0, *∂*^2^ *g / ∂r* ^2^ |_*r*=0_= *e* ^−^ · (1 − *m*^2^) · [−*m* sinh (*r*) + cosh (*r*)]| _*r*=0_ = (1 − *m*^2^) > 0, and hence, *g* _*D*_(*r*) has an absolute minimum at *r* = 0. Thus, *g* _*D*_ (*r*) *g* ≥ (0) = 0. Because *∂g/ ∂r* is strictly positive for *D ∩* {*r* ≠ 0}, *g* _*D ∩ r* ≠ 0_ (*r*) > *g* (0) and thus *g* (*r*) is strictly positive for *r* ≠ 0 and −1 < *m*< 1. Therefore, 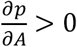 and 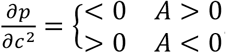.

According to the “false signal” hypothesis, the addition of background odors increases *A*_+_ and *A*_−_, thus increasing *c*^2^, and increasing the probability of a correct response when the target odor is present and decreasing it when it is absent (note that the probability of a correct response is *p* in “target on” trials and 1 − *p* in “target off” trials). Similarly, the “signal reduction” hypothesis posits that background odors decrease *A*_+_ and *A*_−_, with the opposite effects on *p*. Finally, in the “noise boost” hypothesis, background odors increase *c*^2^ without changing *A*_+_ and *A*_−_, which results in a decrease in the probability of a correct response both when the target odor is present and when it is absent.

Next, we consider the effect of odor backgrounds on the speed at which decisions are being made in the “noise boost” mechanism. In the DDM, the mean decision-time,E[*DT*], as a function of the model parameters is given by (Bogacz et al., 2006; Srivastava et al., 2016):

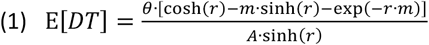

The derivative of E[*DT*] with respect to *c*^2^ is given by

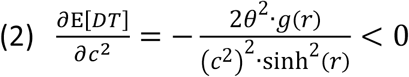

Thus, if background odors increase *c*^2^ then they are predicted to decrease E[*DT*]

## Acknowledgements

This work was funded by a European Research Council Starting Grant (755764—COFBMIX, D.R.), the Israel Science foundation (Grant 757/16, Y.L.), the DFG (CRC 1080, Y.L.), and the Gatsby Charitable Foundation (Y.L.). We thank Mati Joshua and members of the Rokni lab for comments on the manuscript.

## Notes

### Competing Interest Statement

The authors have declared no competing interest.

### Summary of Updates

A clarification about trial duration has been added to the abstract and results sections. Funding sources have been added to the Acknowledgements.

## References

Abraham, N.M., Spors, H., Carleton, A., Margrie, T.W., Kuner, T., and Schaefer, A.T. (2004). Maintaining accuracy at the expense of speed: stimulus similarity defines odor discrimination time in mice. Neuron 44, 865–876.

Araneda, R.C., Kini, A.D., and Firestein, S. (2000). The molecular receptive range of an odorant receptor. Nat Neurosci 3, 1248–1255.

Arevian, A.C., Kapoor, V., and Urban, N.N. (2008). Activity-dependent gating of lateral inhibition in the mouse olfactory bulb. Nat Neurosci 11, 80–87.

Aungst, J.L., Heyward, P.M., Puche, A.C., Karnup, S.V., Hayar, A., Szabo, G., and Shipley, M.T. (2003). Centre–surround inhibition among olfactory bulb glomeruli. Nature 426, 623–629.

Bogacz, R., Brown, E., Moehlis, J., Holmes, P., and Cohen, J.D. (2006). The physics of optimal decision making: a formal analysis of models of performance in two-alternative forced-choice tasks. Psychol Rev 113, 700–765.

Brody, C.D., and Hanks, T.D. (2016). Neural underpinnings of the evidence accumulator. Current Opinion in Neurobiology 37, 149–157.

Crouzet, S.M., and Serre, T. (2011). What are the visual features underlying rapid object recognition? Front. Psychology 2, 326.

Davison, I.G., and Katz, L.C. (2007). Sparse and Selective Odor Coding by Mitral/Tufted Neurons in the Main Olfactory Bulb. J. Neurosci. 27, 2091–2101.

Duchamp-Viret, P., Duchamp, A., and Chaput, M.A. (2003). Single olfactory sensory neurons simultaneously integrate the components of an odour mixture. European Journal of Neuroscience 18, 2690–2696.

Dylla, M., Hrnicek, A., Rice, C., and Ramachandran, R. (2013). Detection of tones and their modification by noise in nonhuman primates. J Assoc Res Otolaryngol 14, 547–560.

Economo, M.N., Hansen, K.R., and Wachowiak, M. (2016). Control of Mitral/Tufted Cell Output by Selective Inhibition among Olfactory Bulb Glomeruli. Neuron 91, 397–411.

Elhilali, M., and Shamma, S.A. (2008). A cocktail party with a cortical twist: How cortical mechanisms contribute to sound segregation. The Journal of the Acoustical Society of America 124, 3751–3771.

Firestein, S., and Shepherd, G.M. (1992). Neurotransmitter antagonists block some odor responses in olfactory receptor neurons. Neuroreport: An International Journal for the Rapid Communication of Research in Neuroscience 3, 661–664.

Fletcher, M.L. (2011). Analytical Processing of Binary Mixture Information by Olfactory Bulb Glomeruli. PLoS ONE 6, e29360.

Franks, K.M., Russo, M.J., Sosulski, D.L., Mulligan, A.A., Siegelbaum, S.A., and Axel, R. (2011). Recurrent Circuitry Dynamically Shapes the Activation of Piriform Cortex. Neuron 72, 49–56.

Giraudet, P., Berthommier, F., and Chaput, M. (2002). Mitral Cell Temporal Response Patterns Evoked by Odor Mixtures in the Rat Olfactory Bulb. J Neurophysiol 88, 829–838.

Gold, J.I., and Shadlen, M.N. (2007). The Neural Basis of Decision Making. Annual Review of Neuroscience 30, 535–574.

Grabska-Barwinska, A., Beck, J., Pouget, A., and Latham, P. (2013). Demixing odors - fast inference in olfaction. In Advances in Neural Information Processing Systems 26, C.J.C. Burges, L. Bottou, M. Welling, Z. Ghahramani, and K.Q. Weinberger, eds. (Curran Associates, Inc.), pp. 1968–1976.

Gupta, P., Albeanu, D.F., and Bhalla, U.S. (2015). Olfactory bulb coding of odors, mixtures and sniffs is a linear sum of odor time profiles. Nat Neurosci 18, 272–281.

Heekeren, H.R., Marrett, S., and Ungerleider, L.G. (2008). The neural systems that mediate human perceptual decision making. Nature Reviews Neuroscience 9, 467–479.

Kadohisa, M., and Wilson, D.A. (2006). Olfactory Cortical Adaptation Facilitates Detection of Odors Against Background. Journal of Neurophysiology 95, 1888–1896.

Kemp, S. (1984). Reaction time to a tone in noise as a function of the signal-to-noise ratio and tone level. Perception & Psychophysics 36, 473–476.

Krajbich, I., and Rangel, A. (2011). Multialternative drift-diffusion model predicts the relationship between visual fixations and choice in value-based decisions. PNAS 108, 13852–13857.

Kurahashi, T., Lowe, G., and Gold, G.H. (1994). Suppression of odorant responses by odorants in olfactory receptor cells. Science 265, 118–120.

Laming, D.R.J. (1968). Information theory of choice-reaction times (Oxford, England: Academic Press).

Lamme, V.A. (1995). The neurophysiology of figure-ground segregation in primary visual cortex. J. Neurosci. 15, 1605–1615.

Lei, H., Mooney, R., and Katz, L.C. (2006). Synaptic integration of olfactory information in mouse anterior olfactory nucleus. J. Neurosci. 26, 12023–12032.

Li, Z., and Hertz, J. (2000). Odour recognition and segmentation by a model olfactory bulb and cortex. Network 11, 83–102.

Malnic, B., Hirono, J., Sato, T., and Buck, L.B. (1999). Combinatorial Receptor Codes for Odors. Cell 96, 713–723.

Mathis, A., Rokni, D., Kapoor, V., Bethge, M., and Murthy, V.N. (2016). Reading Out Olfactory Receptors: Feedforward Circuits Detect Odors in Mixtures without Demixing. Neuron 91, 1110–1123.

McDermott, J.H. (2009). The cocktail party problem. Curr. Biol. 19, R1024–1027.

Meister, M., and Bonhoeffer, T. (2001). Tuning and topography in an odor map on the rat olfactory bulb. J. Neurosci. 21, 1351–1360.

Mendonça, A.G., Drugowitsch, J., Vicente, M.I., DeWitt, E.E.J., Pouget, A., and Mainen, Z.F. (2020). The impact of learning on perceptual decisions and its implication for speed-accuracy tradeoffs. Nature Communications 11, 2757.

Micheyl, C., and Oxenham, A.J. (2010). Pitch, harmonicity and concurrent sound segregation: Psychoacoustical and neurophysiological findings. Hearing Research 266, 36–51.

Najemnik, J., and Geisler, W.S. (2005). Optimal eye movement strategies in visual search. Nature 434, 387–391.

Oka, Y., Omura, M., Kataoka, H., and Touhara, K. (2004). Olfactory receptor antagonism between odorants. The EMBO Journal 23, 120–126.

Penker, S., Licht, T., Hofer, K.T., and Rokni, D. (2020). Mixture coding and segmentation in the anterior piriform cortex. Front. Syst. Neurosci. 14.

Pfister, P., Smith, B.C., Evans, B.J., Brann, J.H., Trimmer, C., Sheikh, M., Arroyave, R., Reddy, G., Jeong, H.-Y., Raps, D.A., et al. (2020). Odorant Receptor Inhibition Is Fundamental to Odor Encoding. Current Biology 30, 2574-2587.e6.

Poo, C., and Isaacson, J.S. (2009). Odor Representations in Olfactory Cortex: “Sparse” Coding, Global Inhibition, and Oscillations. Neuron 62, 850–861.

Poo, C., and Isaacson, J.S. (2011). A Major Role for Intracortical Circuits in the Strength and Tuning of Odor-Evoked Excitation in Olfactory Cortex. Neuron 72, 41–48.

Ratcliff, R. (1978). A theory of memory retrieval. Psychological Review 85, 59–108.

Ratcliff, R., and McKoon, G. (2007). The Diffusion Decision Model: Theory and Data for Two-Choice Decision Tasks. Neural Computation 20, 873–922.

Ratcliff, R., Van Zandt, T., and McKoon, G. (1999). Connectionist and diffusion models of reaction time. Psychol Rev 106, 261–300.

Ratcliff, R., Cherian, A., and Segraves, M. (2003). A Comparison of Macaque Behavior and Superior Colliculus Neuronal Activity to Predictions From Models of Two-Choice Decisions. Journal of Neurophysiology 90, 1392–1407.

Reddy, G., Zak, J.D., Vergassola, M., and Murthy, V.N. (2018). Antagonism in olfactory receptor neurons and its implications for the perception of odor mixtures.

Rinberg, D., Koulakov, A., and Gelperin, A. (2006). Speed-accuracy tradeoff in olfaction. Neuron 51, 351– 358.

Rokni, D., Hemmelder, V., Kapoor, V., and Murthy, V.N. (2014). An olfactory cocktail party: figure-ground segregation of odorants in rodents. Nat Neurosci 17, 1225–1232.

Rospars, J.-P., Lansky, P., Chaput, M., and Duchamp-Viret, P. (2008). Competitive and Noncompetitive Odorant Interactions in the Early Neural Coding of Odorant Mixtures. J. Neurosci. 28, 2659–2666.

Rubin, B.D., and Katz, L.C. (1999). Optical Imaging of Odorant Representations in the Mammalian Olfactory Bulb. Neuron 23, 499–511.

Singh, V., Murphy, N.R., Balasubramanian, V., and Mainland, J.D. (2019). Competitive binding predicts nonlinear responses of olfactory receptors to complex mixtures. PNAS 116, 9598–9603.

Soucy, E.R., Albeanu, D.F., Fantana, A.L., Murthy, V.N., and Meister, M. (2009). Precision and diversity in an odor map on the olfactory bulb. Nat. Neurosci. 12, 210–220.

Srivastava, V., Holmes, P., and Simen, P. (2016). Explicit moments of decision times for single- and double-threshold drift-diffusion processes. Journal of Mathematical Psychology 75, 96–109.

Stettler, D.D., and Axel, R. (2009). Representations of odor in the piriform cortex. Neuron 63, 854–864.

Suzuki, N., and Bekkers, J.M. (2012). Microcircuits Mediating Feedforward and Feedback Synaptic Inhibition in the Piriform Cortex. J. Neurosci. 32, 919–931.

Takeuchi, H., Ishida, H., Hikichi, S., and Kurahashi, T. (2009). Mechanism of olfactory masking in the sensory cilia. The Journal of General Physiology 133, 583–601.

Teki, S., Chait, M., Kumar, S., Kriegstein, K. von, and Griffiths, T.D. (2011). Brain Bases for Auditory Stimulus-Driven Figure–Ground Segregation. J. Neurosci. 31, 164–171.

Uchida, N., and Mainen, Z.F. (2003). Speed and accuracy of olfactory discrimination in the rat. Nature Neuroscience 6, 1224–1229.

Urban, N.N., and Sakmann, B. (2002). Reciprocal intraglomerular excitation and intra- and interglomerular lateral inhibition between mouse olfactory bulb mitral cells. The Journal of Physiology 542, 355–367.

Wilson, D.A. (2003). Rapid, Experience-Induced Enhancement in Odorant Discrimination by Anterior Piriform Cortex Neurons. Journal of Neurophysiology 90, 65–72.

Wolfson, S.S., and Landy, M.S. (1998). Examining edge- and region-based texture analysis mechanisms. Vision Res. 38, 439–446.

Xu, L., Li, W., Voleti, V., Zou, D.-J., Hillman, E.M.C., and Firestein, S. (2020). Widespread receptor-driven modulation in peripheral olfactory coding. Science 368.

Yoshida, I., and Mori, K. (2007). Odorant Category Profile Selectivity of Olfactory Cortex Neurons. J. Neurosci. 27, 9105–9114.

Zak, J.D., Reddy, G., Vergassola, M., and Murthy, V.N. (2020). Antagonistic odor interactions in olfactory sensory neurons are widespread in freely breathing mice. Nature Communications 11, 3350.

Zariwala, H.A., Kepecs, A., Uchida, N., Hirokawa, J., and Mainen, Z.F. (2013). The limits of deliberation in a perceptual decision task. Neuron 78, 339–351.

